# Tissue-specificity of gene expression diverges slowly between orthologs, and rapidly between paralogs

**DOI:** 10.1101/065086

**Authors:** Nadezda Kryuchkova-Mostacci, Marc Robinson-Rechavi

**Author notes:** Author of Correspondence: Marc Robinson-Rechavi, Department of Ecology and Evolution, University of Lausanne, Switzerland.

## Abstract

The ortholog conjecture implies that functional similarity between orthologous genes is higher than between paralogs. It has been supported using levels of expression and Gene Ontology term analysis, although the evidence was rather weak and there were also conflicting reports. In this study on 12 species we provide strong evidence of high conservation in tissue-specificity between orthologs, in contrast to low conservation between within-species paralogs. This allows us to shed a new light on the evolution of gene expression patterns. While there have been several studies of the correlation of expression between species, little is known about the evolution of tissue-specificity itself. Ortholog tissue-specificity is strongly conserved between all tetrapod species, with the lowest Pearson correlation between mouse and frog at r = 0.66. Tissue-specificity correlation decreases strongly with divergence time. Paralogs in human show much lower conservation, even for recent Primate-specific paralogs. When both paralogs from ancient whole genome duplication tissue-specific paralogs are tissue-specific, it is often to different tissues, while other tissue-specific paralogs are mostly specific to the same tissue. The same patterns are observed using human or mouse as focal species, and are robust to choices of datasets and of thresholds. Our results support the following model of evolution: in the absence of duplication, tissue-specificity evolves slowly, and tissue-specific genes do not change their main tissue of expression; after small-scale duplication the less expressed paralog loses the ancestral specificity, leading to an immediate difference between paralogs; over time, both paralogs become more broadly expressed, but remain poorly correlated. Finally, there is a small number of paralog pairs which stay tissue-specific with the same main tissue of expression, for at least 300 million years.

**Author summary:** From specific examples, it has been assumed by comparative biologists that the same gene in different species has the same function, whereas duplication of a gene inside one species to create several copies allows them to acquire different functions. Yet this model was little tested until recently, and then has proven harder than expected to confirm. One of the problems is defining “function” in a way which can be easily studied. We introduce a new way of considering function: how specific is the activity (“expression”) of a gene? Genes which are specific to certain tissues have functions related to these tissues, whereas genes which are broadly active over many or all tissues have more general functions for the organism. We find that this “tissue-specificity” evolves very slowly in the absence of duplication, while immediately after duplication the new gene copy differs. This shows that indeed duplication leads to a strong increase in the evolution of new functions.

## Introduction

The ortholog conjecture is widely used to transfer annotation among genes, for example in newly sequenced genomes. But has been difficult to establish whether and how much orthologs share more similar functions than paralogs [1,2]. The most widely accepted model is that orthologs diverge slower, and that the generation of paralogs through duplication leads to strong divergence and even change of function. It is also expected that in general homologs diverge functionally with time. The test of these hypotheses poses fundamental questions of molecular evolution, about the rate of functional evolution and the role of duplications, and is essential to the use of homologs in genome annotations.

Surprisingly, there are several studies which have reported no difference between orthologs and paralogs, or even the opposite, that paralogs would be more functionally similar than orthologs. Tests of the ortholog conjecture using sequence evolution found no difference after speciation or duplication in positive selection [3], nor in amino acid shifts [4]. The debate was truly launched by Nehrt et al. [5] who reported in a large scale study, based on expression levels similarity and Gene Ontology (GO) analysis in human and mouse, that paralogs are better predictors of function than orthologs. Of note, methodological aspects of the GO analysis of that study were criticized by several other authors [6,7]. Using a very similar GO analysis but correcting biases in the data, from 13 bacterial and eukaryotic species, Altenhoff et al. [8] found more functional similarity between orthologs than between paralogs based on GO annotation analysis, but the differences were very slight.

An early comparison of expression profiles of orthologs in human and mouse reported that they were very different, close to paralogs and even to random pairs [9]. Further studies, following Nehrt et al. [5], found little or no evidence for the ortholog conjecture in expression data. Rogozin et al. [10] reported that orthologs are more similar than between species paralogs but less similar than within-species paralogs based on correlations between RNA-seq expression profiles in human and mouse. Wu et al. [11] found only a small difference between orthologs and paralogs. Paralogs were significantly more functionally similar than orthologs, but by classifying in subtypes they reported that one-to-one orthologs are the most functionally similar. The analysis was done on the level of function by looking at expression network similarities in human, mouse, fly and worm.

On the other hand, the ortholog conjecture has been supported by several studies of gene expression. *Contra* Yanai et al. [9], several studies have reported good correlations between expression levels of orthologs, between human and mouse [12], or among amniotes [13]. Moreover, some studies have reported changes of expression following duplication, although without explicitly testing for the ortholog conjecture: duplicated genes are more likely to show changes in expression profiles than single-copy genes [14,15]. Chung et al. [16] reported through network analysis in human that duplicated genes diverge rapidly in their expression profile. Recently Assis and Bachtrog [17] reported that paralog function diverges rapidly in mammals. They analysed among other things difference in tissue-specificity between a pair of paralogs and their single copy ortholog in closely related species. They conclude that divergence of paralogs results in increased tissue-specificity, and that there are differences between tissues. Finally, several explicit tests of the ortholog conjecture have also found support using expression data. Huerta-Cepas et al. [18] reported that paralogs have higher levels of expression divergence than orthologs of the similar age, using microarray data with calls of expressed/not expressed in human and mouse. They also claimed that a significant part of this divergence was acquired shortly after the duplication event. Chen and Zhang [7] re-analysed the RNA-seq dataset of Brawand et al. [13] and reported that expression profiles of orthologs are significantly more similar than within-species paralogs. Thus while the balance of evidence appears to weight towards confirmation of the ortholog conjecture, functional data has failed so far to strongly support or invalidate it. Even results which support the ortholog conjecture often do so with quite slight differences between orthologs and paralogs [8,10]. Yet expression data especially should have the potential to solve this issue, since it provides functional evidence for many genes in the same way across species, without the ascertainment biases of GO annotations or other collections of small scale data. Part of the problem is that the relation between levels of expression and gene function is not direct, making it unclear what biological signal is being compared in correlations of these levels. Another problem is that the comparison of different transcriptome datasets between species suffers from biases introduced by ubiquitous genes [19] or batch effects [20].

In our analysis we have concentrated on the tissue-specificity of expression. Tissue-specificity indicates in how many tissues a gene is expressed, and whether it has large differences of expression level between them. It reflects the functionality of the gene: if the gene is expressed in many tissues then it is “house keeping” and has a function needed in many organs and cell types; tissue-specific genes have more specific roles, and tissue adjusted functions. Recent results indicate that tissue-specificity is conserved between human and mouse orthologs, and that it is functionally informative [21]. Moreover, tissue-specificity can be computed in a comparable manner in different animal datasets without notable biases, as long as at least 6 tissues are represented, including preferably testis, nervous system, and proportionally not too many parts of the same organ (e.g. not many parts of the brain).

Are there major differences between the evolution of tissue-specificity after duplication (paralogs) or without duplication (orthologs)? We analyse the conservation of one-to-one orthologs and within-species paralogs with evolutionary time, using RNA-seq datasets from 12 species.

## Results

We compared orthologs between 12 species: human, chimpanzee, gorilla, macaque, mouse, rat, cow, opossum, platypus, chicken, frog, and fruit fly. Overall 7 different RNA-seq datasets were used, including 6 to 27 tissues (see Materials and Methods). Three comparisons were performed with the largest sets as focal data: 27 human tissues from Fagerberg et al., 16 human tissues from Bodymap, and 22 tissues from mouse ENCODE [22–24]. For all analyses we used tissue-specificity of expression as described in Materials and Methods.

The first notable result is that tissue-specificity is strongly correlated between one-to-one orthologs. The correlations between human and four other species are presented in Fig 1a for illustration. This confirms and extends our previous observation [21], which was based on one human and one mouse datasets. Correlation of tissue-specificity varies between 0.74 and 0.89 among tetrapods, and is still 0.43 between human and fly, 0.38 between mouse and fly. The latter is despite the very large differences in anatomy and tissue sampling between the species compared, showing how conserved tissue-specificity can be in evolution.

The correlation between orthologs decreases with divergence time (Fig 2). The decline is linear. An exponential model is not significantly better: ANOVA was not significantly better for the model with log_10_ of time than for untransformed time for any dataset *(p >* 0.0137, *q >* 1%). The trend is not caused by the outlier fly data point: removing it there is still a significant decrease of correlation for orthologs (see Supplementary Materials). Results are also robust to the use of Spearman instead of Pearson correlation between tissue-specificity values.

**Fig 1.**
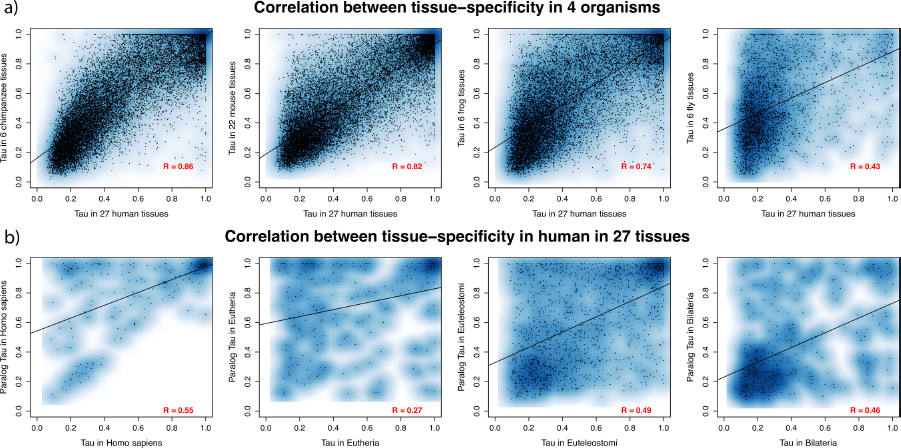
Pearson correlation of tissue-specificity between a) orthologs and b) paralogs. a) Human ortholog vs. one-to-one ortholog in another species; b) highest expressed paralog vs. lowest expressed paralog in human, for different duplication dates.

The correlation between within-species paralogs is significantly lower than between orthologs (ANOVA *p*<0.0137, *q*<1% for all datasets) (Fig 2). Moreover, there is no significant decline in correlation with evolutionary time (neither linear nor exponential) for paralogs. This may indicate almost immediate divergence of paralogs upon duplication, although other scenarios are possible (see Discussion).

The results are consistent using human or mouse as focal species (Fig 2a and b). Results are also consistent using a different human RNA-seq dataset (Fig S1).

**Fig 2.**
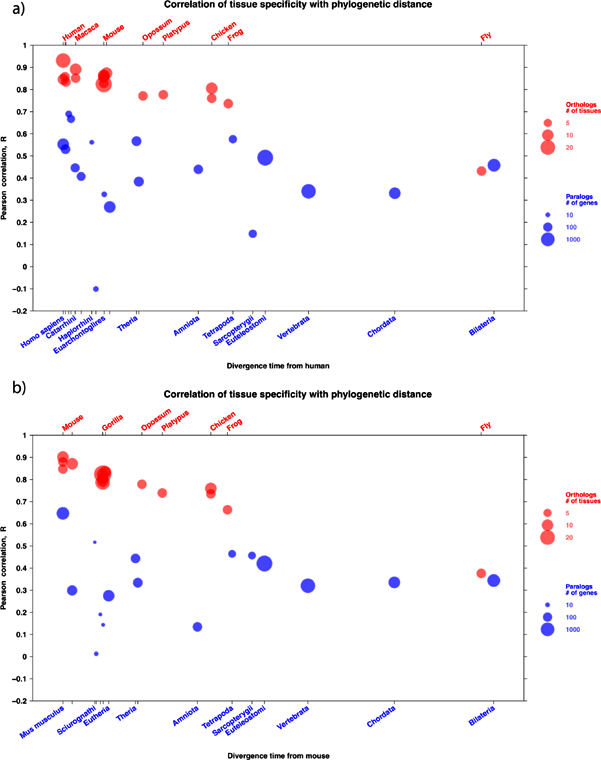
Pearson correlation of tissue-specificity focusing on a) human and b) mouse. X-axis, divergence time in million years between the genes compared; Y-axis, Pearson correlation between values of τ over genes. In red, the correlation of orthologs between the focal species and other species; representative species are noted above the figure; there are several points when there are several datasets for a same species, e.g. four for mouse (Table 1); the size of red circles is proportional to the number of tissues used for calculation of tissue-specificity. In blue, the correlation of paralogs in the focal species, according to the date of duplication; representative taxonomic groups for this dating are noted under the figure; the size of blue circles is proportional to the number of genes in the paralog group.

This main analysis is based on the correlation of tissue-specificity for orthologs called pairwise between species. The number of orthologs used in the analysis is thus variable (available in Supplementary Materials). An additional analysis was also performed using the same orthologs for all tetrapods, 4785 genes (Fig S2-S4). Correlations of these “conserved orthologs” are not significantly different from those observed over all orthologs.

The analysis was also performed on all the datasets with tissue-specificity calculated without testis (Fig S5-S7). The correlation between orthologs becomes significantly lower (ANOVA *p*=0.000178), while between paralogs it does not change significantly (ANOVA *p*=0.846). Even though the correlation between orthologs becomes weaker there is still a significant difference between orthologs and paralogs (ANOVA *p*=1.299e-07). The same analysis was also performed removing 4 other main tissues (brain, heart, kidney and liver) (Fig S8–S11). For the brain the correlation between orthologs becomes significantly lower (ANOVA *p*=0.000289), but stays higher than for paralogs; for other tissues there is no significant difference. For paralogs the correlation never changes significantly.

We also performed the analysis removing genes on sex chromosomes (Fig S12–S14). This analysis was done without frog, as sex chromosome information is not available. This does not change significantly the correlations between either orthologs (ANOVA *p*=0.856) or paralogs (ANOVA *p*=0.755).

In general paralogs have lower expression and are more tissue-specific than orthologs (Fig S15), which is consistent with the dosage-sharing model [25,26]. Young paralogs are very tissue-specific, and get more ubiquitous with divergence time (Fig 1b and Fig S16); this is true for all datasets, and for τ calculated with or without testis. We also tested for asymmetry by comparing paralog pairs to the closed possible non duplicated outgroup; e.g., we compared each Eutheria specific paralog to the non duplicated opossum outgroup (one-to-two ortholog; Fig 3). We observe that the higher expressed paralog has a stronger correlation with the outgroup, thus appears to keep more the ancestral tissue-specificity, while the lower expressed paralog has a lower correlation and appears to become more tissue-specific (Fig 3), which is consistent with a form of neo-functionalization.

**Fig 3.**
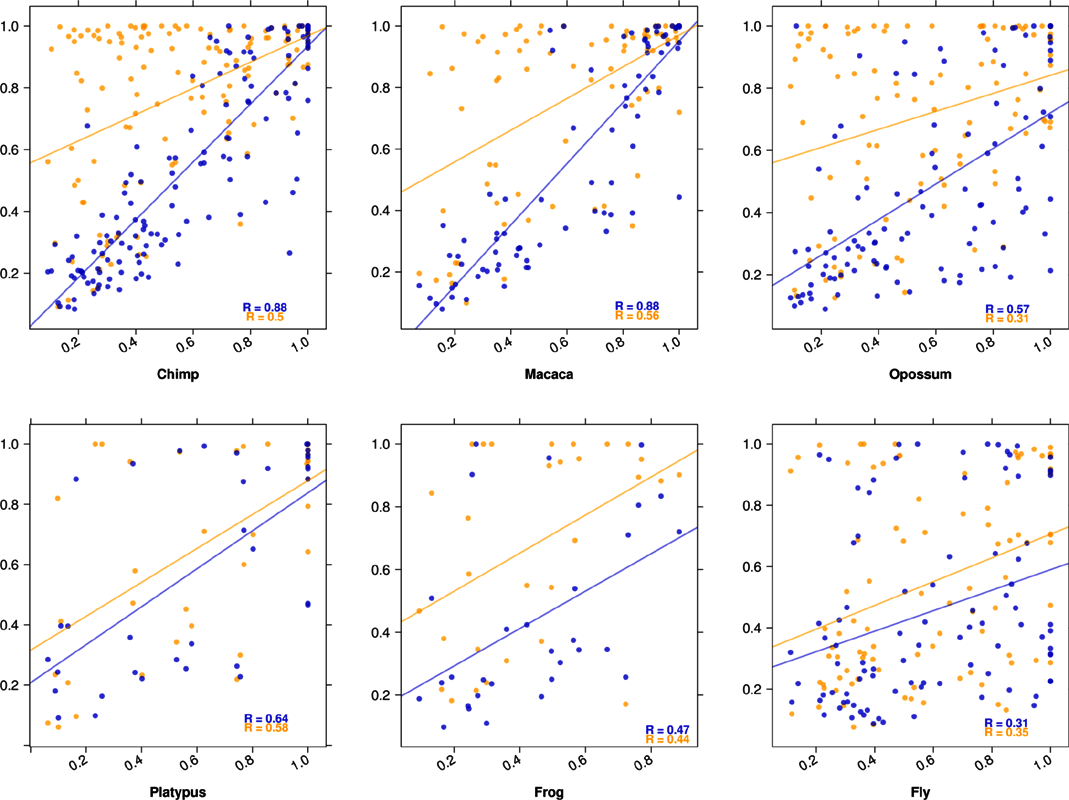
Distribution of tissue-specificity in paralogs compared to an outgroup ortholog. For each graph, paralogs of a given phylogenetic age are compared to the closest outgroup un-duplicated ortholog; thus these paralogs are “in-paralogs” relative to the speciation node, and are both “co-orthologs” to the outgroup. X-axis, τ of unduplicated ortholog. Y-axis, τ of paralogs. Blue points are values for the paralog with highest maximal expression of the pair of paralogs, orange points are values for the other.

When both orthologs of a pair are tissue-specific (τ > 0.8), they are most often expressed in the same tissue (Fig 4). The same is observed when both paralogs are tissue-specific and are younger than the divergence of tetrapods. But for Euteleostomi and Vertebrata paralogs, if both are tissue-specific then they are as likely to be expressed in the different as in same tissues; most of these are expected to be ohnologs, i.e. due to whole genome duplication. This analysis was performed on the Brawand et al. (2011) dataset, because it has the most organisms with the same 6 tissues. This result does not change after removing testis (Fig S17), nor changing the τ threshold from 0.8 to 0.3 (Fig S18–S19). Also after removing all tissue-specific genes (τ > 0.8), the difference between orthologs and paralogs is smaller but stay significant (ANOVA *p*=0.001) (Fig S20).

**Fig 4.**
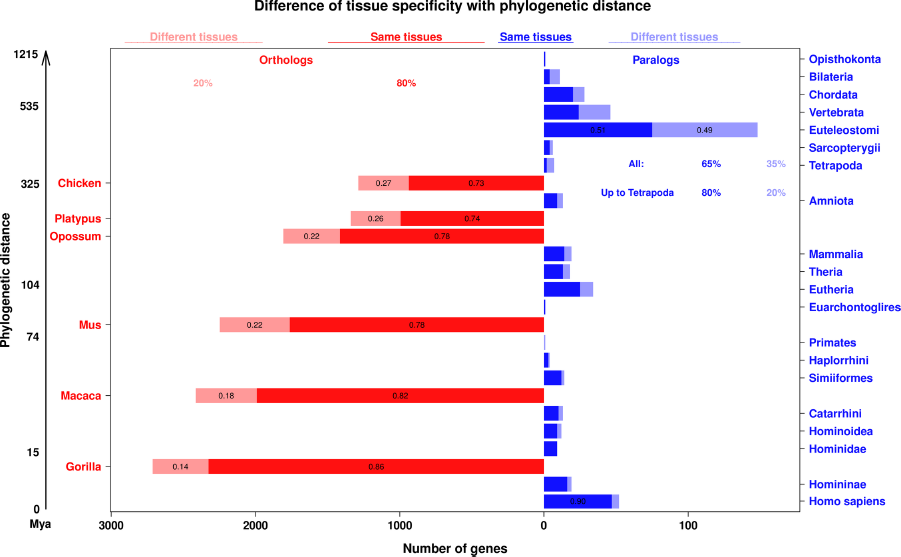
Difference of tissue-specificity between orthologs and paralogs. Each bar represents the number of gene pairs of a given type for a given phylogenetic age, for which both genes of the pair are tissue-specific (τ > 0.8). In dark color, the number of gene pairs specific to the same tissue; in light color, the number of gene pairs specific to different tissues. Orthologs are in red, in the left panel, paralogs are in blue, on the right panel; notice that the scales are different for orthologs and for paralogs. Orthologs are one-to-one orthologs to human and paralogs are within-species paralogs in human. The overall proportions of pairs in the same or different tissues are indicated for orthologs and paralogs; in addition, for paralogs the proportion for pairs younger than the divergence of tetrapods (whole genome duplication) is also indicated.

## Discussion

Our results show that most genes have their tissue-specificity conserved between species. This provides strong new evidence for the evolutionary conservation of expression patterns. Using tissue-specificity instead of expression values allows easy comparison between species, as bias of normalisation or use of different datasets has little effect on results [21]. All of our results were confirmed using three different focus datasets, from human or mouse, and thus appear to be quite robust.

The conservation of expression tissue-specificity of protein coding genes that we find is high even for quite distant one-to-one orthologs: the Pearson correlation between τ in human or mouse and τ in frog is R = 0.74 (respectively R = 0.66) over 361 My of divergence. Even between fly and mammals it is more than 0.38. Moreover, this tissue-specificity can be easily compared over large datasets without picking a restricted set of homologous tissues (e.g. in [7,13]). The correlation between orthologs is strongest for recent speciations, and decreases linearly with divergence time. This decrease shows that we are able to detect a strong evolutionary signal in tissue-specificity, which has not always been obvious in functional comparisons of orthologs (e.g. [5,8]).

Correlation between within-species paralogs is much lower than between orthologs. Whereas the expression of young paralogs has been recently reported to be highly conserved [17], we find a large difference between even very young paralogs in tissue-specificity. In Assis and Bachtrog [17], the measure of tissue-specificity is not clearly defined, but it seems to be TSI [27], which performed poorly as an evolutionarily relevant measure in our recent benchmark [21]; they also treated female and male samples as different “tissues”, confounding two potentially different effects. The low correlation that we observed for young paralogs does not decrease significantly with divergence time. It is possible that on the one hand paralogs do diverge in tissue-specificity with time, and that on the other hand this trend is compensated by biased loss of the most divergent paralogs. It is also possible that we lack statistical power to detect a slight decrease in correlation of paralogs, due to low numbers of paralogs for many branches of the phylogeny. The most likely interpretation is that for small-scale paralogs (defined as not from whole genome duplication [28]) there is an asymmetry, with a daughter gene which lacks regulatory elements of the parent gene upon birth; further independent changes in tissue-specificity in each paralog would preserve the original lack of correlation. In any case, we do not find support for a progressive divergence of tissue-specificity for paralogs.

The overall conservation of tissue-specificity could be due to a subset of genes, and most notably sex-related genes. Indeed, the largest set of tissue-specific genes are testis-specific [21]. To verify the influence of sex-related genes, we performed all analyses without testis expression data, or without genes mapped to sex chromosomes. After removing testis expression from all datasets the correlation between paralogs does not change significantly, while between orthologs is gets significantly weaker. The lower correlation of orthologs suggests that testis specific genes are conserved between species, and as they constitute a high proportion of tissue-specific genes, they contribute strongly to the correlation. Removing sex chromosome located genes does not change results significantly. After removing testis expression the differences of conservation of tissue-specificity between orthologs and paralogs stay significant. Overall, it appears that tissue-specificity calculated with testis represents a true biological signal, and given its large effect it is important to include this tissue in analyses.

In general paralogs are more tissue-specific and have lower expression levels. This could be explained if ubiquitous genes are less prone to duplication or duplicate retention. Yet we do not observe any bias in the orthologs of duplicates towards more tissue-specific genes (Fig 3; see also Supplementary Materials). With time both paralogs get more broadly expressed (Fig 1 and Fig S16). In the rare case where both paralogs are tissue-specific, small-scale young paralogs are expressed in the same tissue, while genome-wide old paralogs (ohnologs) are expressed in different tissues (Fig 4). With the data available, we cannot distinguish the effects of paralog age and of duplication mechanism, since many old paralogs are due to whole genome duplication in vertebrates, whereas that is not the case for the young paralogs. In many cases the higher expressed paralog has a similar tissue-specificity to the ancestral state, while the lower expressed paralog is more tissue-specific (Fig 3).

We have studied gene specificity without taking in account alternative splicing, or the possibility that different transcripts are expressed in different tissues, because it is still difficult to call transcript level expression reliably [29]. This would probably not change our main observations, that tissue-specificity is conserved among orthologs, diverges with evolutionary time, and follows the ortholog conjecture. Of note, recent results have not supported an important role of alternative splicing for differences in transcription between tissues [30,31].

The overall picture that we obtain for the evolution of tissue-specificity is the following. In the absence of duplication, tissue-specificity evolves slowly, thus is mostly conserved, and tissue-specific genes do not change their main tissue of expression (Fig 2 and 4). After small-scale duplication (i.e., not whole genome) paralogs diverge rapidly in tissue-specificity, or already differ at birth. This difference is mostly due to the less expressed paralog losing the ancestral specificity, while the most expressed paralog keeps at first closer to the ancestral state, as estimated from a non duplicated outgroup ortholog (Fig 3). But over time, even the most expressed paralog diverges much more strongly than a non duplicated ortholog. While paralog divergence is rapid, in the small number of genes which stay tissue-specific for both paralogs the main tissue of expression is mostly conserved, for several hundred million years (i.e. origin of tetrapods, Fig 4). With increasing age of the paralogs, they both tend to become more broadly expressed (Fig 1 and Fig S16) while keeping a low correlation. For whole genome duplicates we have less information, because of the age of the event in vertebrates and the lack of good outgroup data. The main difference is that when two genome duplication paralogs are both tissue-specific, they are often expressed in different tissues (Fig 4).

We have used tissue-specificity to estimate the conservation of function, rather than Gene Ontology annotations or expression levels. We believe that this metric is less prone to systematic errors, whether annotation biases for the Gene Ontology, or proper normalisation between datasets and choice of few tissues for expression levels. Our results confirm the Ortholog Conjecture on data which is genome-wide and functionally relevant: orthologs are more similar than within-species paralogs. Moreover, orthologs diverge monotonically with time, as expected. On the contrary, even young paralogs show large differences.

## Material and Methods

RNA-seq data from 12 species (human, gorilla, chimpanzee, macaque, mouse, platypus, opossum, chicken, gorilla, cow, frog, rat and fruit fly) were used for the analysis. We recovered all animal RNA-seq data sets which cover at least 6 adult tissues, and were either pre-processed in Bgee [32], or provided pre-processed data from the publication, as of June 2015. For human, mouse and chicken we used several datasets. All the datasets with the corresponding number of tissues are summarized in Table 1. The numbers of genes used for the analysis are in Table S1 and S2.

The orthology and paralogy calls and their phylogenetic dating for paralogs were taken from Ensembl Compara (Version 75) [33]. Phylogenetic dating was converted to absolute dates using the TimeTree data base [34].

**Table 1.** Datasets used in the paper.

| **Organisms/ datasets** | Fagerberg | Brawand | Bodymap | ENCODE | Necsulea | Merkin | Keane |
| --- | --- | --- | --- | --- | --- | --- | --- |
| **Dataset ID** | E-MTAB-1733 | GSE30352 | GSE30611 | GSE36025 (mouse) | GSE43520 | GSE41637 | GSE30617 |
| **RPKM/FPKM source** | Supp. mat. | Bgee | Bgee | Supp. mat., [35] | Bgee | Bgee | Bgee |
| Human<br>*Homo sapiens* | 27 | 8 | 16 |  |  |  |  |
| Gorilla<br>*Gorilla gorilla* |  | 6 |  |  |  |  |  |
| Chimpanzee<br>*Pan troglodytes* |  | 6 |  |  |  |  |  |
| Macaque<br>*Macaca mulatta* |  | 6 |  |  |  | 9 |  |
| Mouse<br>*Mus musculus* |  | 6 |  | 22 |  | 9 | 6 |
| Rat<br>*Rattus norvegicus* |  |  |  |  |  | 9 |  |
| Cow<br>*Bos taurus* |  |  |  |  |  | 9 |  |
| Opossum<br>*Monodelphis domestica* |  | 6 |  |  |  |  |  |
| Platypus<br>*Ornithorhynchus anatinus* |  | 6 |  |  |  |  |  |
| Chicken<br>*Gallus gallus* |  | 6 |  |  |  | 9 |  |
| Frog<br>*Xenopus tropicalis* |  |  |  |  | 6 |  |  |
| Fly<br>*Drosophila melanogaster* |  |  |  | 6 |  |  |  |
| **Citations** | [22] | [13] | [23] | [24,36] | [37] | [38] | [39] |

For the human dataset from Fagerberg et al. [22] and the fly dataset [36], FPKM values were downloaded from the respective papers Supplementary Materials; the mouse ENCODE project dataset was processed by an in house script (TopHat and Cufflinks [40]); all other data were processed by the Bgee pipeline [32]. For all analyses gene models from Ensembl version 75 were used [41]. Only protein-coding genes were used for analysis. For the analysis of paralogs the youngest couple was taken (Fig S21), and sorted according to the maximal expression, i.e. the reference paralog (called “gene” in our R scripts) is always the one with the highest maximal expression. This choice gives the highest correlation compared to a random sorting (Fig S22).

Analyses were performed in R version 3.2.1 [42] using Lattice [43], plyr [44], gplots [45] and qvalue [46,47] libraries.

As a measure for tissue-specificity we used τ (Tau) [48]:

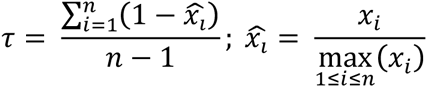

Tau is calculated on the log RNA-seq expression data. The values of τ vary from 0 to 1, where 0 means ubiquitous expressed genes and 1 specific genes. We have recently shown that τ is the best choice for calculating tissue specificity among existing methods [21]. For comparing tissue-specific genes, they were called with τ ≥ 0.8, and assigned to the tissue with the highest expression.

A special case is testis-specificity, as many more genes are expressed in testis than other tissues. For control analysis, all genes with maximal expression in testis were called “testis specific”, independently of τ value.

Over all ANOVA tests performed (112 tests), we used a q-value threshold of 1% of false positives, corresponding to a p-value threshold of 0.066.

## Acknowledgements

We thank Julien Roux and Jialin Liu for their helpful comments and suggestions.

## Supplementary Materials

Supplementary Materials are available online.

## Supplementary Materials

**Additional Supplementary files are available on Figshare:**

https://figshare.com/articles/Tissue-specificity_of_gene_expression_diverges_slowly_between_orthologs_and_rapidly_between_paralogs/3493010

**Table S1.** Number of protein coding genes used for the analysis.

| Organisms/data sets | Fagerberg | Brawand | Bodymap | ENCODE | Necsulea | Merkin | Keane |
| --- | --- | --- | --- | --- | --- | --- | --- |
| Human | 18569 | 19151 | 19113 |  |  |  |  |
| Gorilla |  | 17069 |  |  |  |  |  |
| Chimp |  | 16507 |  |  |  |  |  |
| Macaca |  | 18297 |  |  |  | 19749 |  |
| Mouse |  | 18086 |  | 19442 |  | 18538 | 16892 |
| Rat |  |  |  |  |  | 19215 |  |
| Cow |  |  |  |  |  | 17634 |  |
| Opossum |  | 16622 |  |  |  |  |  |
| Platypus |  | 19036 |  |  |  |  |  |
| Chicken |  | 14332 |  |  |  | 14780 |  |
| Frog |  |  |  |  | 15499 |  |  |
| Fly |  |  |  | 10960 |  |  |  |

**Table S2.** Number of one-to-one orthologous genes of organisms to human, used for the main analysis.

| Organisms/data sets | Fagerberg | Brawand | Bodymap | ENCODE | Necsulea | Merkin | Keane |
| --- | --- | --- | --- | --- | --- | --- | --- |
| Human | - | 17170 | 17224 |  |  |  |  |
| Gorilla |  | 14813 |  |  |  |  |  |
| Chimp |  | 15282 |  |  |  |  |  |
| Macaca |  | 14578 |  |  |  | 14943 |  |
| Mouse |  | 14397 |  | 14876 |  | 14791 | 14056 |
| Rat |  |  |  |  |  | 14040 |  |
| Cow |  |  |  |  |  | 14666 |  |
| Opossum |  | 12445 |  |  |  |  |  |
| Platypus |  | 10490 |  |  |  |  |  |
| Chicken |  | 11352 |  |  |  | 11525 |  |
| Frog |  |  |  |  | 11462 |  |  |
| Fly |  |  |  | 2750 |  |  |  |

**Legend for figures:**

Fig S1 — Fig S4 and Fig S6 — Fig S11: X-axis, divergence time in million years between the genes compared; Y-axis, Pearson correlation between values of τ over genes. In red, the correlation of orthologs between the focal species and other species; representative species are noted above the figure; there are several points when there are several datasets for a same species; the size of red circles is proportional to the number of tissues used for calculation of tissue specificity. In blue, the correlation of paralogs in the focal species, according to the date of duplication; representative taxonomic groups for this dating are noted under the figure; the size of blue circles is proportional to the number of genes in the paralog group.

Fig S14 — Fig S16: Each bar represents the number of gene pairs of a given type for a given phylogenetic age, for which both genes of the pair are tissue-specific. In dark colour, the number of gene pairs specific of the same tissue; in light colour, the number of gene pairs specific of different tissues. Orthologs are in red, in the left panel, paralogs are in blue, on the right panel; notice that the scales are different for orthologs and for paralogs. The overall proportions of pairs in the same or different tissues are indicated for orthologs and paralogs; in addition, for paralogs the proportion for pairs younger than the divergence of tetrapods is also indicated.

**Fig S1.**
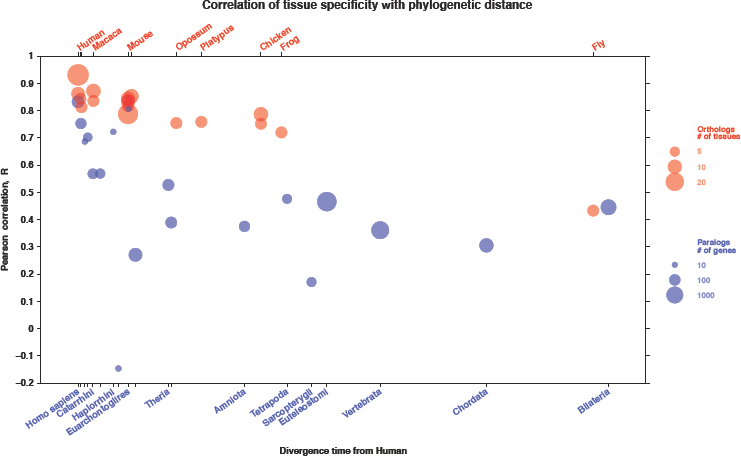
Pearson correlation of tissue specificity according to human Bodymap dataset.

**Fig S2.**
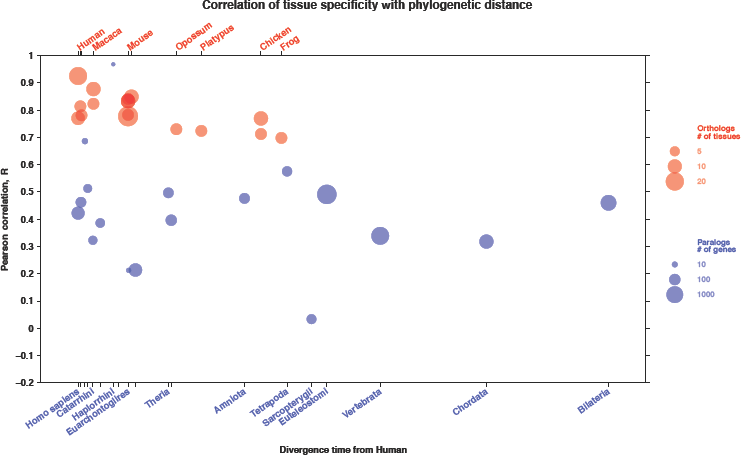
Pearson correlation of tissue specificity according to human Fageberg dataset. Only conserved orthologs (up to frog, present in all analysed species).

**Fig S3.**
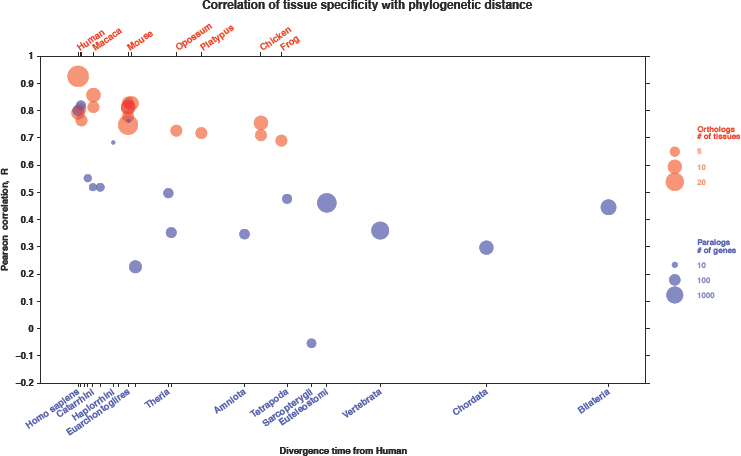
Pearson correlation of tissue specificity according to human Bodymap dataset. Only conserved orthologs (up to frog, present in all analysed species).

**Fig S4.**
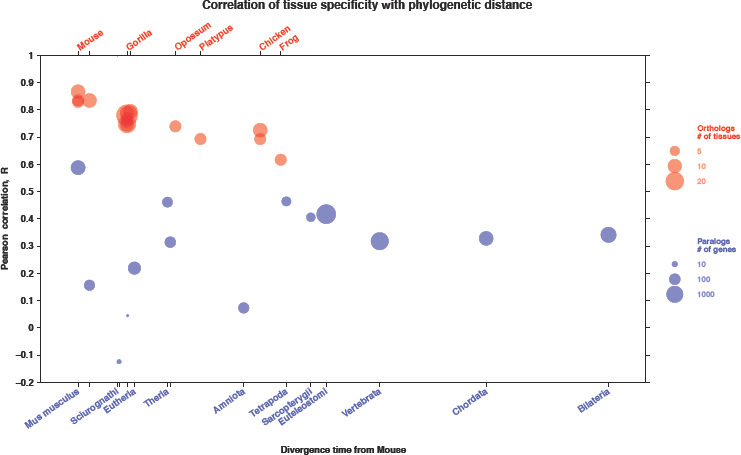
Pearson correlation of tissue specificity according to mouse dataset. Only conserved orthologs (up to frog, present in all analysed species).

**Fig S5.**
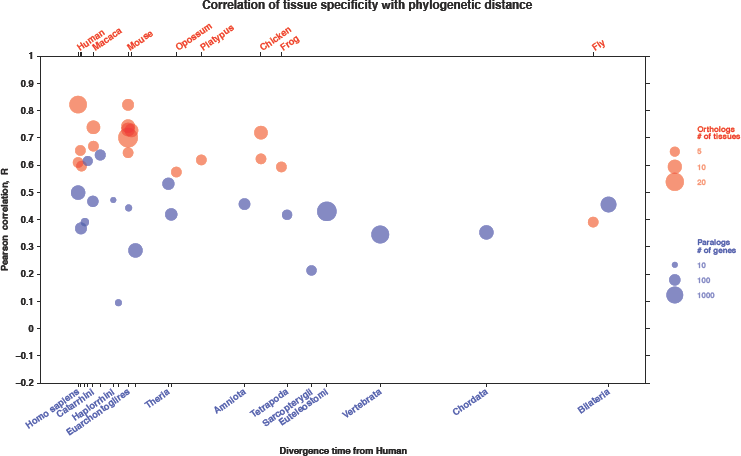
Pearson correlation of tissue specificity according to human Fagerberg dataset. Tissue-specificity calculated without testis.

**Fig S6.**
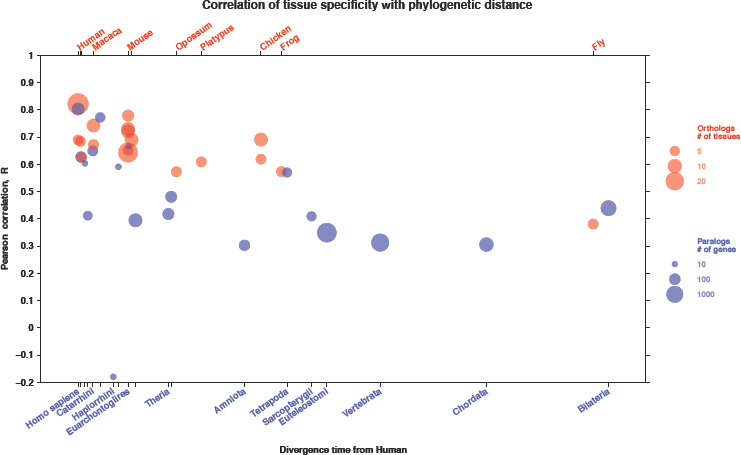
Pearson correlation of tissue specificity according to human Bodymap dataset. Tissue-specificity calculated without testis.

**Fig S7.**
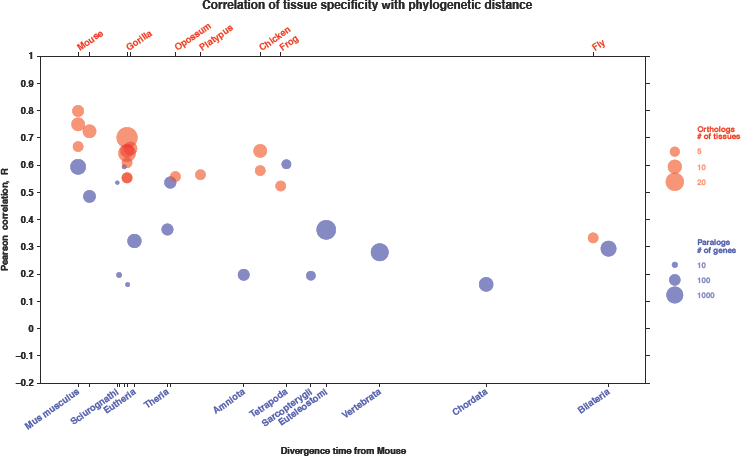
Pearson correlation of tissue specificity according to mouse dataset. Tissue-specificity calculated without testis.

**Fig S8.**
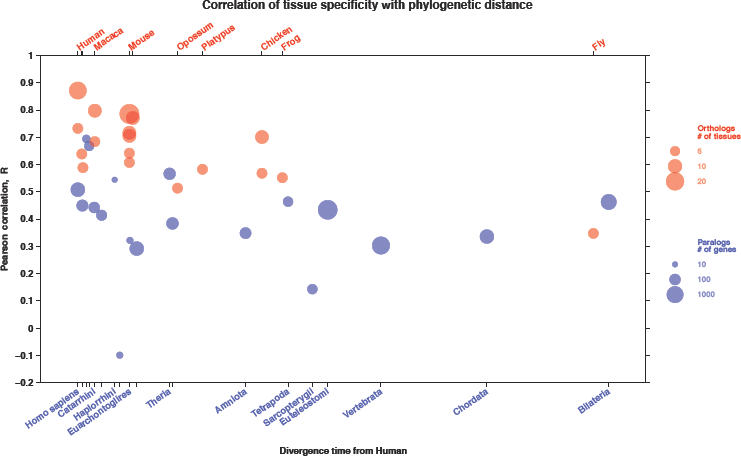
Pearson correlation of tissue specificity according to mouse dataset. Tissue-specificity calculated without brain.

**Fig S9.**
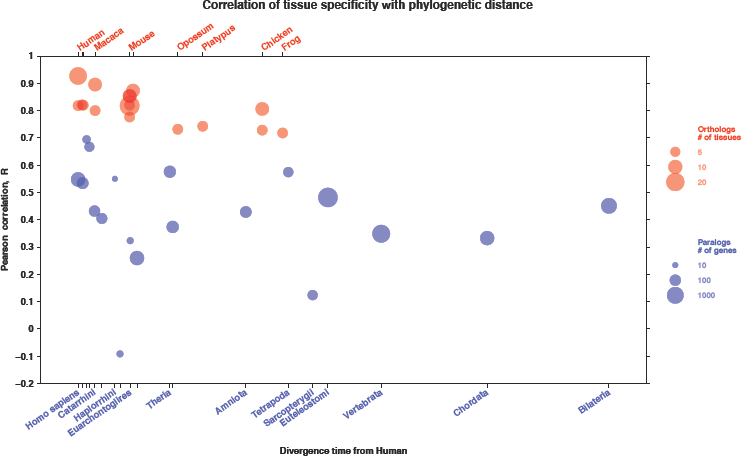
Pearson correlation of tissue specificity according to mouse dataset. Tissue-specificity calculated without heart.

**Fig S10.**
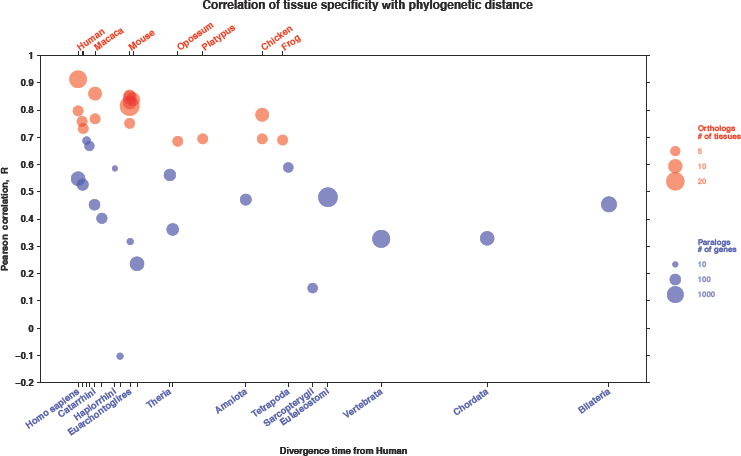
Pearson correlation of tissue specificity according to mouse dataset. Tissue-specificity calculated without kidney.

**Fig S11.**
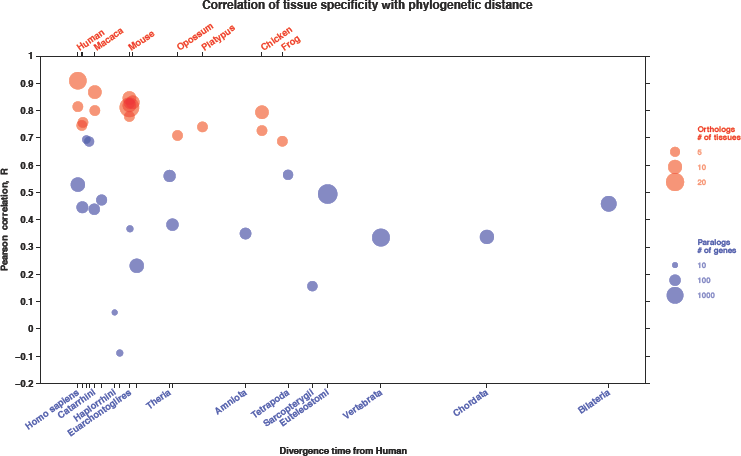
Pearson correlation of tissue specificity according to mouse dataset. Tissue-specificity calculated without liver.

**Fig S12.**
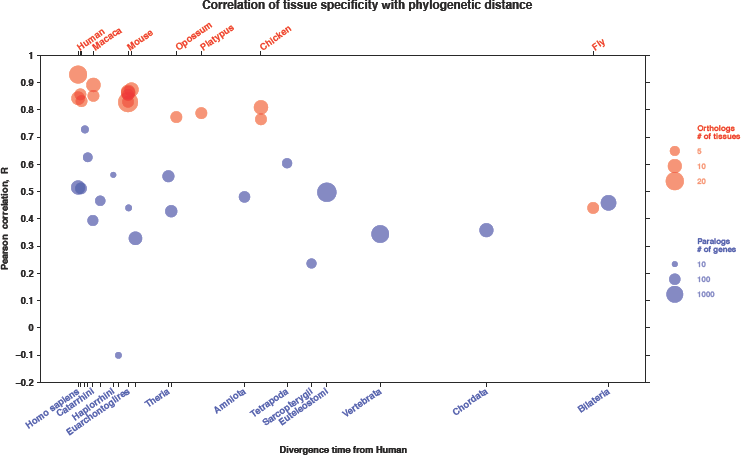
Pearson correlation of tissue specificity according to human Fagerberg dataset. Tissue-specificity calculated without sex-chromosome genes.

**Fig S13.**
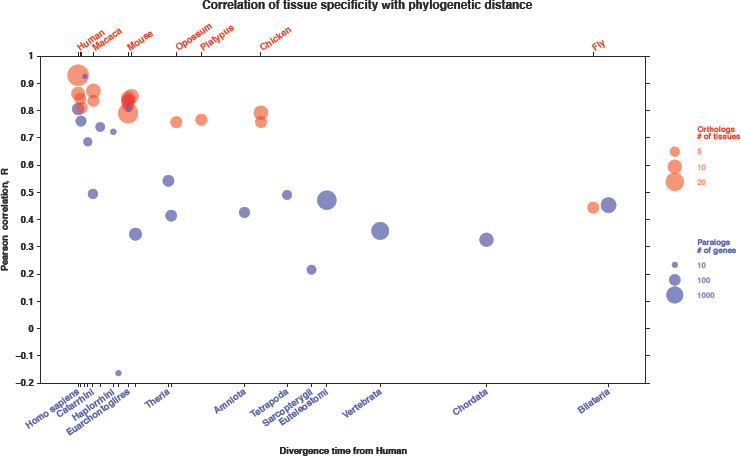
Pearson correlation of tissue specificity according to human Bodymap dataset. Tissue-specificity calculated without sex-chromosome genes.

**Fig S14.**
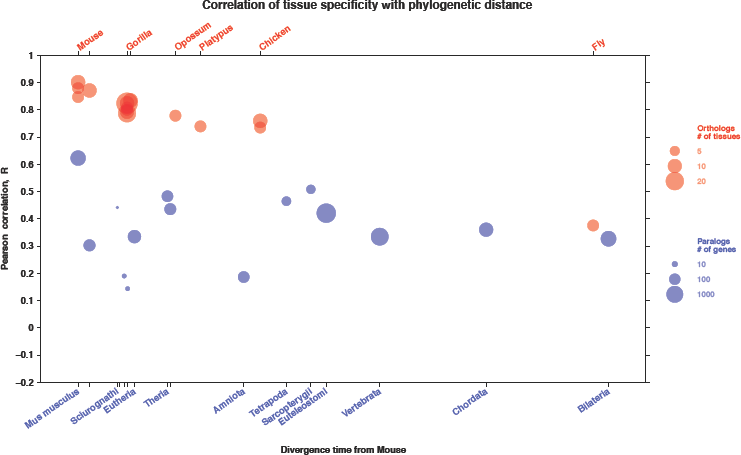
Pearson correlation of tissue specificity according to mouse dataset. Tissue-specificity calculated without sex-chromosome genes.

**Fig S15.**
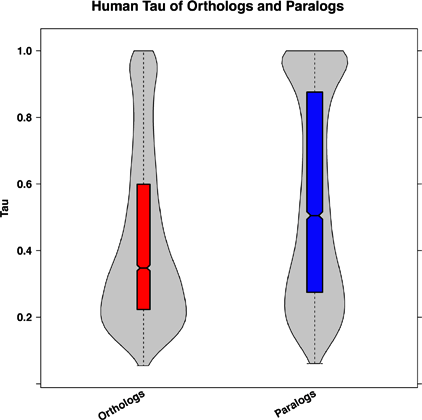
Distribution of tissue-specificity between orthologs and paralogs.

**Fig S16.**
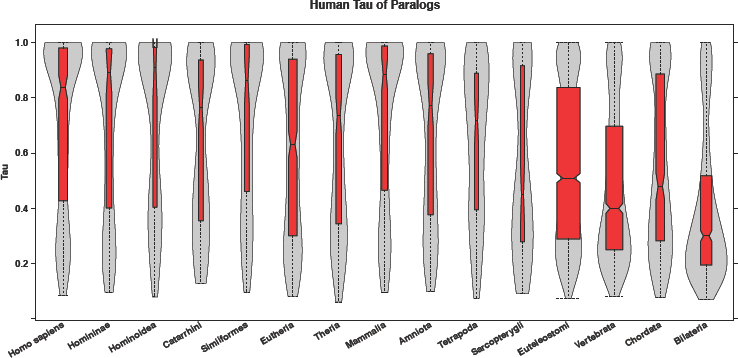
Distribution of tissue-specificity in paralogs of different age of duplication.

**Fig S17.**
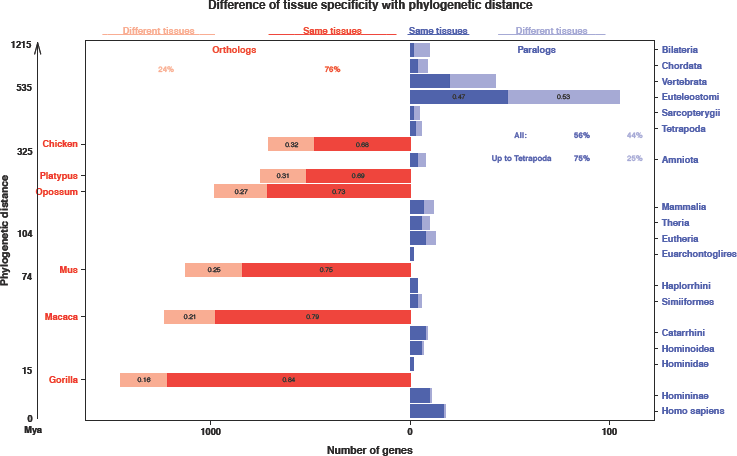
Difference of tissue-specificity between orthologs and paralogs. Tau cut-off 0.8 and calculated without testis.

**Fig S18.**
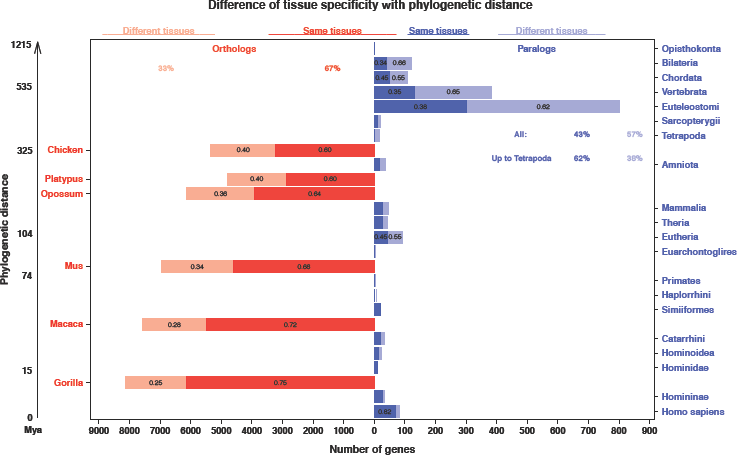
Difference of tissue-specificity between orthologs and paralogs. Tau cut-off 0.3.

**Fig S19.**
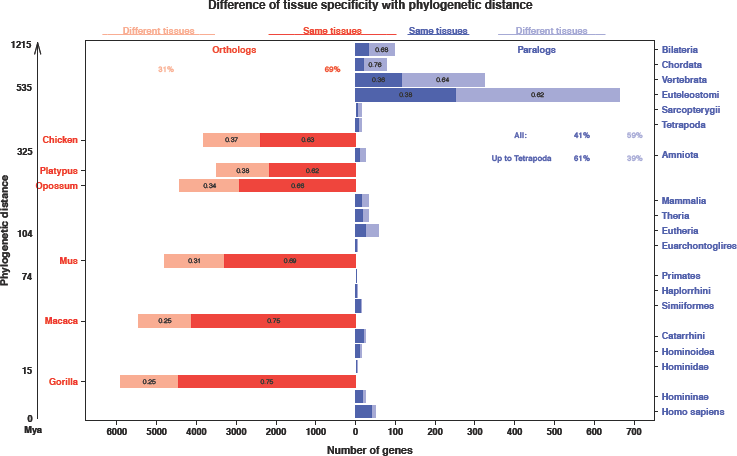
Difference of tissue-specificity between orthologs and paralogs. Tau cut-off 0.3 and calculated without testis.

**Fig S20.**
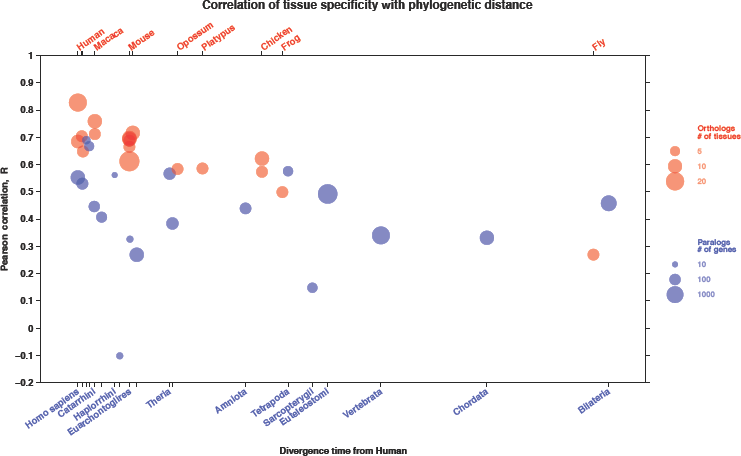
Pearson correlation of tissue specificity according to mouse dataset. Tissue-specificity calculated without tissue-specific genes (Tau > 0.8).

**Fig S21.**
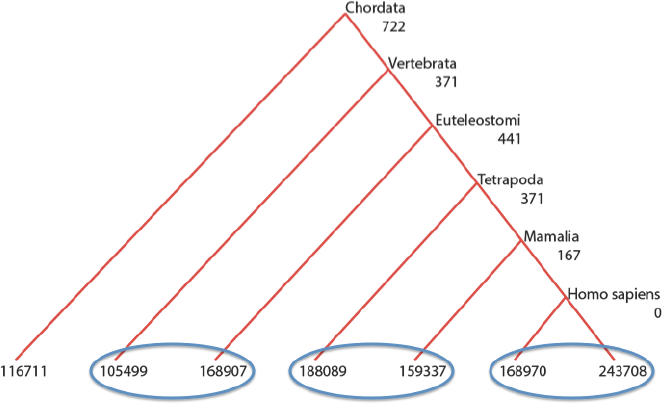
Choice of paralogs. The tree is the example for one paralog family. The blue circles represent how the youngest couple of paralogs was chosen for different phylogenetic ages. Gene names are of the form ENSG00000xxxxxx, with xxxxxx to be replaced with the numbers shown on the figure.

**Fig S22.**
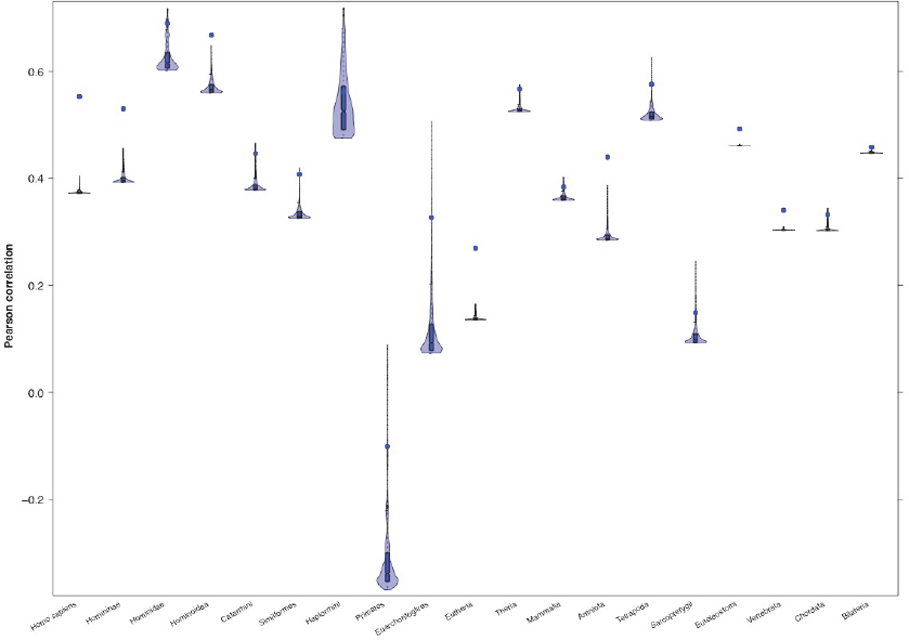
Pearson correlations between paralogs. Box plots represents 1000 random attribution of paralogs in each pair to the x and y vectors for the correlation. The blue dot is the correlation between the paralogs sorted as in the main analysis, i.e. the highest expressed in x and the lowest in y for each pair.

